# Characterisation of cefotaxime-resistant urinary *Escherichia coli* from primary care in South-West England 2017-2018

**DOI:** 10.1101/701383

**Authors:** Jacqueline Findlay, Virginia C. Gould, Paul North, Karen E. Bowker, O. Martin Williams, Alasdair P. MacGowan, Matthew B. Avison

**Affiliations:** School of Cellular & Molecular Medicine, Biomedical Sciences Building, University of Bristol, University Walk, Bristol, UK; Department of Infection Sciences, Severn Infection Partnership, Southmead Hospital, Bristol, United Kingdom

## Abstract

**Objectives:** Third-generation cephalosporin-resistant *Escherichia coli* from community-acquired urinary tract infections (UTI) have been increasingly reported worldwide. In this study we sought to determine and characterise the mechanisms of cefotaxime-resistance (CTX-R) employed by urinary *E. coli* obtained from primary care over a 12-month period, in Bristol and surrounding counties in the South West of England.

**Methods:** Cephalexin resistant (Ceph-R) *E. coli* isolates were identified directly from general practice (GP) referred urine samples using disc susceptibility testing as per standard diagnostic procedures. CTX-R was determined by subsequent plating onto MIC breakpoint plates. β-Lactamase genes were detected by PCR. Whole Genome Sequencing (WGS) was performed on 225 urinary isolates and analyses were performed using the Centre for Genomic Epidemiology platform. Patient information provided by the referring GPs was reviewed.

**Results:** During the study period, Ceph-R *E. coli* (n=900) were obtained directly from urines from 146 GPs. Seventy-percent (626/900) of isolates were CTX-R. WGS of 225 non-duplicate isolates identified that the most common mechanism of CTX-R was *bla*_CTX-M_ carriage (185/225; 82.2%), predominantly *bla*_CTX-M-15_ (114/185; 61.6%), followed by carriage of plasmid mediated AmpCs (pAmpCs) (17/225; 7.6%), ESBL *bla*_SHV_ variants (6/225; 2.7%), AmpC hyperproduction (13/225; 5.8%), or a combination of both *bla*_CTX-M_ and pAmpC carriage (4/225; 1.8%). Forty-four sequence types (STs) were identified with ST131 representing 101/225 (45.0%) of sequenced isolates, within which the *bla*_CTX-M-15_-positive clade C2 was dominant (54/101; 53.5%). Ciprofloxacin-resistance (CIP-R) was observed in 128/225 (56.9%) of sequenced CTX-R isolates – predominantly associated with fluoroquinolone-resistant clones ST131 and ST1193.

**Conclusions:** Most Ceph-R urinary *E. coli*s were CTX-R, predominantly caused by *bla*_CTX-M_ carriage. There was a clear correlation between CTX-R and CIP-R, largely attributable to the dominance of the high-risk pandemic clones, ST131 and ST1193 in this study. This localised epidemiological data provides greater resolution than regional data and can be valuable for informing treatment choices in the primary care setting.

## Introduction

*Escherichia coli* that are resistant to β-lactam antibiotics, particularly to cephalosporins, represent a major global public health concern. Third-generation cephalosporins (3GCs), such as cefotaxime, are used across the world to treat infections caused by *E. coli* (e.g. urinary tract [UTIs], bloodstream and intra-abdominal infections) and subsequently the emergence of resistance is particularly worrisome.^1^ Resistance to 3GCs in *E. coli* can be caused by multiple mechanisms including chromosomally encoded AmpC β-lactamase hyperproduction, and may involve increased efflux, and reduced outer membrane permeability, but is predominantly attributed to the spread of plasmid-mediated AmpC (pAmpC, e.g. *bla*_CMY_) or extended-spectrum β-lactamases (ESBLs, e.g. *bla*_CTX-M_).^2^ *E. coli* which harbour ESBLs are often co-resistant to multiple antibiotic classes and subsequently the treatment options for such infections may be limited.^3^

UTIs are the most common bacterial infection type in both primary and hospital care settings in the developed world,^4^ and are associated with considerable morbidity.^5^ Previous studies of community-acquired UTIs in several mainland European countries found that *E. coli* were the most commonly isolated uropathogen, accounting for over half of all isolates (53.3-76.7%).^6-8^ The incidence of community-acquired UTI in the UK is difficult to determine since such infections are not reportable and most are diagnosed and treated in a primary care setting, with diagnosis often based solely upon patient symptoms rather than a positive urine culture. Community-acquired UTIs are most often treated empirically and subsequently local epidemiological data is useful for informing treatment choice. Treatment failure for community-acquired UTIs, particularly in immune-compromised patients, increases the risk of the infection spreading to other sites including the bloodstream, with grave consequences.^9, 10^

*E. coli* sequence types (STs) belonging to phylogroups B2 (STs 73, 95 and 131) and D (ST69) have been reported to be major causes of both UTIs and bloodstream infections (BSIs) in the UK.^11^ Since its initial description in 2008, numerous studies have shown that the multidrug-resistant pandemic clone, ST131, is a major cause of UTI globally.^12-14^ The ST131 clonal group can be broken down by population genetics into three clades based on their association with particular *fimH* types: A/*fimH*41, B/*fimH*22, and C/*fimH*30.^15^ Clade C can be further broken down into four subclades; C1 – not usually associated with ESBL carriage but typically fluoroquinolone resistant (FQ-R), C1-M27 and C1-nM27 – both associated with *bla*_CTX-M-27_ carriage and FQ-R, and C2 (also known as H30Rx) – associated with *bla*_CTX-M-15_ carriage and FQ-R.^15, 16^ Studies have suggested that the global dominance of ESBL-positive ST131 is, in part, due to its increased virulence potential over its non-ST131 ESBL-positive counterparts.^17, 18^

This study sought to use whole genome sequencing (WGS) to characterise the population structure and determine the mechanisms of CTX-R employed by urinary *E. coli* isolates referred from general practice in Bristol and surrounding counties in the South-West of England serving a population of approximately 1.2 million people.

## Materials and Methods

### Bacterial isolates, identification and susceptibility testing

Cephalexin-resistant (Ceph-R) urinary *E. coli* isolates were obtained from routine urine microbiology at Severn Infection Partnership Southmead Hospital. Urine samples were submitted between Sept 2017 and Aug 2018, from 146 general practices located throughout Bristol and including coverage in Gloucestershire, Somerset and Wiltshire.

Bacterial identification was carried out using BD™ CHROMagar™ Orientation Medium chromogenic agar (BD, GmbH, Heidelberg, Germany).

Antibiotic susceptibilities were performed by disc testing or, in the case of colistin, by broth microdilution and interpreted according to EUCAST guidelines.^19^ Ceph-R isolates were subcultured onto agar plates containing 2 mg/L cefotaxime (CTX) and isolates that were positive for growth were deemed CTX-resistant (CTX-R), and taken forward for further testing.

### Screening for β-lactamase genes

Two multiplex PCRs were performed to screen for β-lactamase genes. The first to detect *bla*_CTX-M_ groups as previously described,^20^ and the second to detect the following additional β-lactamase genes; *bla*_CMY-2_ type, *bla*_DHA_, *bla*_SHV_, *bla*_TEM_, *bla*_OXA-1_, using the primers listed in Table S1.

### WGS and analyses

WGS was performed by MicrobesNG (https://microbesng.uk/) on a HiSeq 2500 instrument (Illumina, San Diego, CA, USA) using 2×250 bp paired end reads. Reads were trimmed using Trimmomatic,^21^ assembled into contigs using SPAdes^22^ 3.13.0 (http://cab.spbu.ru/software/spades/) and contigs were annotated using Prokka.^23^ Resistance genes, plasmid replicon types, sequence types and *fim* types were assigned using the ResFinder,^24^ PlasmidFinder,^25^ MLST^26^ 2.0 and FimTyper on the Center for Genomic Epidemiology (http://www.genomicepidemiology.org/) platform.

ST131 clades were identified by resistance gene carriage and *fimH* type, and in the case of clade C1-M27, by the presence of the prophage region M27PP1 (CP006632)^16^ through sequence alignment using progressive Mauve alignment software.^27^

MLST and resistance gene data were analysed to produce a minimum spanning tree using Bionumerics software v7.6 (Applied Maths, Sint-Martens-Latem, Belgium).

Plasmid pUB_DHA-1 was sequenced to closure and submitted to GenBank with accession number MK048477. Reads were mapped using Geneious Prime 2019.1.3 (https://www.geneious.com).

### Plasmid Transformation

Transformation of plasmid extractions from isolates encoding *mcr-1* and *bla*_OXA-244_ were attempted by electroporation using *E. coli* DH5 alpha as a recipient. Transformants were selected, respectively, on LB agar containing 0.5 mg/L colistin, or containing 100 mg/L ampicillin with a 10 µg ertapenem disc being placed on the agar surface (Oxoid Ltd, Basingstoke, UK). Transformants were confirmed by PCR (Table S1).

### Analysis of patient demographic information

Limited, non-identifiable patient information was obtained from the request forms sent with submissions from referring GP practices.

## Results and Discussion

### Patient demographics and antimicrobial susceptibilities

Nine hundred Ceph-R urinary *E. coli* isolates, obtained from primary care in Bristol and surrounding areas, were collected during the period of the study. Most isolates were obtained from female patients (721/900; 80.1%) and the average patient age was 63.2 years. Seventy-percent (626/900) were CTX-R and within these, most were again from females (507/626; 81.0%) with an average patient age of 63.6 years.

### β-Lactamase genes of interest detected by PCR in CTX-R isolates

Table 1 indicates the number of CTX-R isolates carrying each β-lactamase genes of interest (GOIs; *bla*_CTX-Ms_, *bla*_CMY_, *bla*_DHA_ or *bla*_SHV_). *bla*_CTX-Ms_ were by far the prevalent gene group, found in 571/626 (91.2%) of isolates. Within these *bla*_CTX-M-G1_ were most common (421/626) followed by *bla*_CTX-M-G9_ (149/626) and *bla*_CTX-M-G8_ (one isolate). pAmpCs *bla*_CMY_ and *bla*_DHA_ were found in 13 (3 alongside *bla*_CTX-M-G1_) and 17 (4 alongside *bla*_CTX-M-G1_) isolates respectively. *bla*_SHV_ was found in 11 (3 alongside *bla*_CTX-M-G1_) isolates and the remaining 24 isolates harboured none of the GOIs as detected by PCR.

**Table 1.**
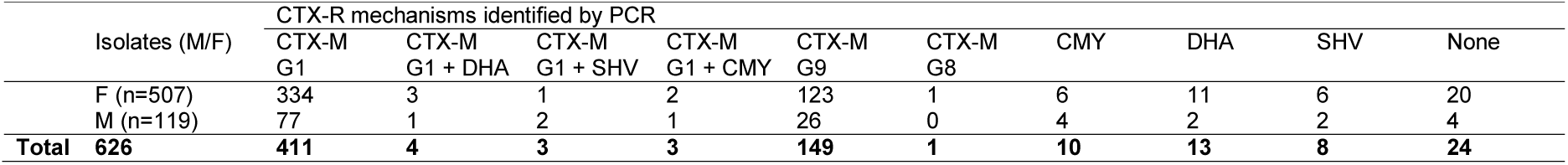
Beta-lactamase genes detected by multiplex PCRs on 626 CTX-R isolates.

### Whole Genome Sequencing (WGS) analyses

#### GOI variants and STs

Two-hundred and twenty-five isolates, chosen to be representative of resistance gene carriage (as previously determined by PCR) and patient demographics (age, sex) obtained throughout the entire study period, were selected for WGS. Within these, 44 sequence types (STs) were identified, with numbers of isolates ranging from 1 to 101 representatives. ST131 was dominant (n=101), followed by STs 69 (n=15), 73 (n=15), 38 (n=13), 1193 (n=11), and 10 (n=8). The remaining 38 STs had 1 to 4 representative isolates (Figure 1).

**Figure 1.**
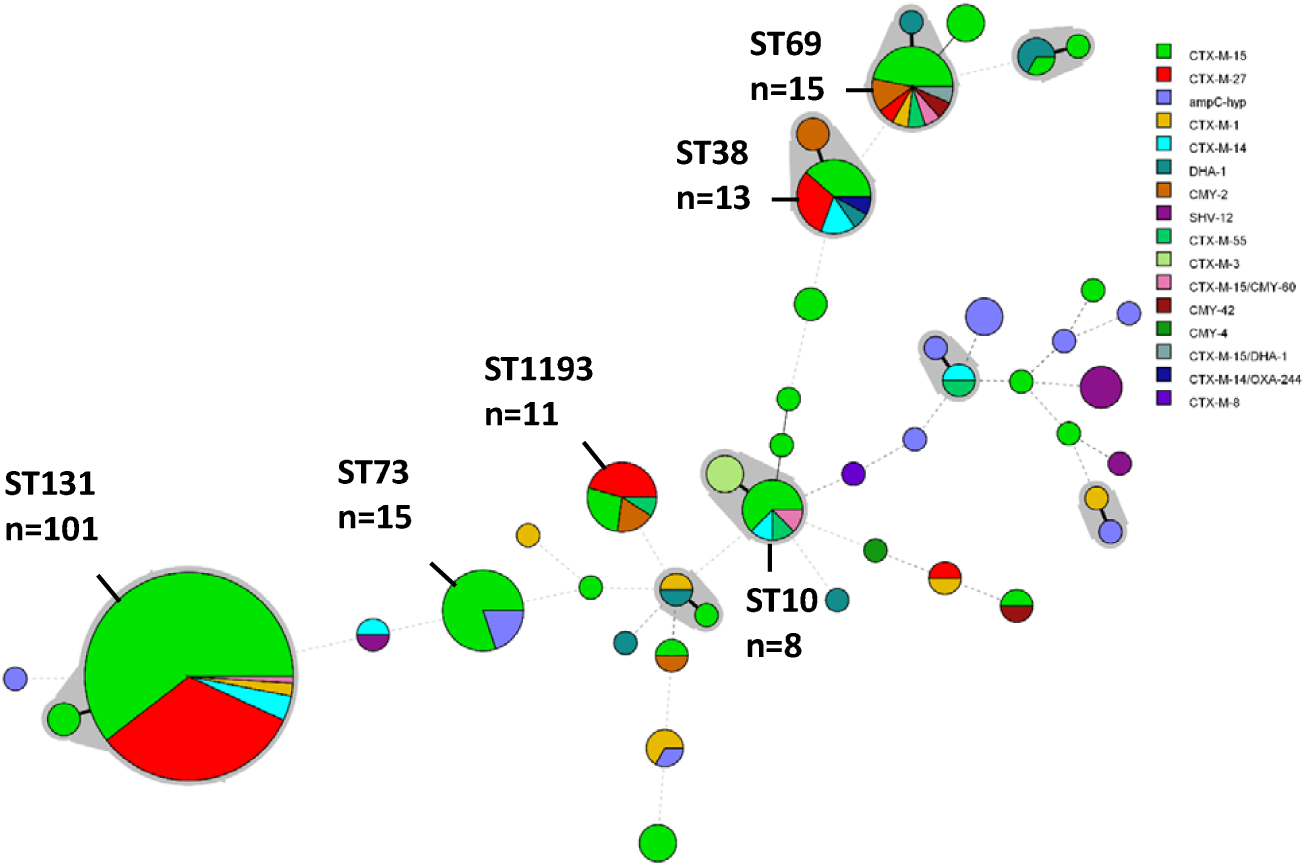
Minimum spanning tree of the MLST profiles of 225 CTX-R *E. coli* isolates. The shaded areas represent single locus variants (SLVs). Members of the most prevalent STs (>4 representatives) are labelled and their number of representatives indicated. The diameter of the circle represents the number of isolates of that particular ST and the coloured segments indicate which CTX-R mechanisms were identified. Thick solid lines represent SLVs, thin solid lines represent double-locus variants and dashed connecting lines indicate multilocus variants.

CTX-R GOIs were identified in all but thirteen isolates (212/225; 94.2%). Eighty-four percent (189/225) of isolates harboured one of seven *bla*_CTX-M_ gene variants (Table 2). Carriage of *bla*_CTX-M-15_ was the most common CTX-R mechanism identified (118/189) followed by *bla*_CTX-M-27_ (44/189) and *bla*_CTX-M-14_ (10/189). Amongst the non-CTX-M GOIs, four *bla*_CMY_ variants were identified; *bla*_CMY-2_ (n=7), *bla*_CMY-4_ (n=1), *bla*_CMY-42_ (n=2) and *bla*_CMY-60_ (n=3; all three being co-carried alongside *bla*_CTX-M-15_), as well as *bla*_DHA-1_ (n=8; one alongside *bla*_CTX-M-15_) and *bla*_SHV-12_ (n=6). The narrow spectrum β-lactamases *bla*_OXA-1_, *bla*_TEM-1_ and inhibitor-resistant variant *bla*_TEM-33_ were found in 53, 82, and one isolate respectively.

**Table 2.** ESBL/pAmpC variants identified in the 225 isolates subjected to WGS.

| *bla*<sub>CTX-M</sub> variant |  |  |  |  |  |  | *bla*<sub>CMY</sub> variant |  |  |  | *bla*<sub>DHA</sub><br>variant | *bla*<sub>SHV</sub><br>variant | None<br>detected |
| --- | --- | --- | --- | --- | --- | --- | --- | --- | --- | --- | --- | --- | --- |
| CTX-M-1 | CTX-M-3 | CTX-M-8 | CTX-M-14 | CTX-M-15 | CTX-M-27 | CTX-M-55 | CMY-2 | CMY-4 | CMY-42 | CMY-60 | DHA-1 | SHV-12 | NA |
| 9 | 3 | 1 | 10 | 118 | 44 | 4 | 7 | 1 | 2 | 3 | 8 | 6 | 13 |

Note: 3 isolates harboured both *bla*<sub>CMY-60</sub> and *bla*<sub>CTX-M-15</sub>, and one isolate harboured *bla*<sub>DHA-1</sub> and *bla*<sub>CTX-M-15</sub>.

#### AmpC-hyperproducing isolates

All thirteen (5.8%) sequenced isolates, where no CTX-R GOI could be identified, were presumed to be chromosomal AmpC β-lactamase hyperproducers because they carry mutations within the *ampC* promoter/attenuator region previously seen in confirmed AmpC hyperproducers (Table 3).^28, 29^ These represented nine different STs, each having one representative, with the exceptions of the STs 75 and 200 of which there were three representatives each (Table S2). This indicates a lack of dominant clones in AmpC hyperproducers identified in this study. If we go on to assume that the 24 isolates negative for GOIs by PCR are AmpC-hyperproducers, as was found with the thirteen representative sequenced isolates, then 3.8% of the isolates in this study could be classed as AmpC-hyperproducers. Additionally, one *bla*_CTX-M-15_-positive isolate was found to also harbour *ampC* promoter changes associated with hyperproduction.

**Table 3.** Mutations found within promoter/attenuator region of the 13 presumed AmpC-hyperproducing isolates subjected to WGS relative to *E. coli* MG1655 (Genbank Accession Number NC_000913.3).

| No. of isolates | *ampC* promoter/attenuator mutations | Pribnow box |
| --- | --- | --- |
| 8 | -42C>T, -25G>A, -1C>T, +57C>T | TTGACA - 17nt - TATCGT |
| 2 | -28G>A, ins -12 T -13, +22G>T | TTGTCA - 17nt - TACAAT |
| 2 | -11C>T, ins -12 T -13, +33G>A, +36G>A | TTGTCA - 17nt - TATAAT |
| 1 | -32T>A, +34C>A, +57C>T | TTGACA - 16nt - TACAAT |

#### Characterisation of ST131 Isolates

##### ESBLs and clades

One hundred and one ST131 isolates harboured the following CTX-R mechanisms/alleles; *bla*_CTX-M-1_ (n=1), *bla*_CTX-M-14_ (n=4), *bla*_CTX-M-15_ (n=61), *bla*_CTX-M-15_/*bla*_CTX-M-60_ (n=1), *bla*CTX-M-27 (n=33). The isolates were broken down into their respective clades (Table 4); ST131-C2 was dominant (54/101; 53.5%) followed by C1-M27 (M27) (23/101; 22.8%), A (11/101; 10.9%), C1-nM27 (5/101; 5.0%), one clade B, and seven isolates were unclassified. Eighty-eight percent (89/101) of isolates, and notably all clade C2 isolates, were CIP-R as is a typical characteristic of this lineage. Two non-ST131 members of the ST131 complex, both of which were *bla*_CTX-M-15_-positive ST8313 isolates (a *fumC* single locus variant (SLV) of ST131), also harboured the same chromosomal FQ-R associated mutations in *gyrA, parC* and *parE* as are associated with ST131/C2 – suggesting this ST may be ST131/C2 derived. Previous studies have highlighted the dominance of ST131 and particularly the clade C2/H30Rx on a worldwide scale.^13, 14^ Since its initial description in 2008 in isolates from 3 continents,^12, 13^ ST131 has been reported across all inhabited continents.^15^ The recently described C1 subclade, C1-M27, as characterised by the presence of a 11,894 bp prophage-like genomic island M27PP1, was initially described in Japan in 2016 and has been reported in countries in at least three continents: Europe, Asia and North America, so far.^16^ The presence of C1-M27 isolates in this study indicates the expansion of this particular ST131 sublineage into the UK, similarly to that which has been reported from countries in mainland Europe.^30^

**Table 4.** ST131 clades and CTX-R GOI alleles harboured by 101 isolates subject to WGS.

|  |  |  | CTX-R Alleles |  |  |  |  |
| --- | --- | --- | --- | --- | --- | --- | --- |
| ST131 Clades (no.) |  |  | CTX-M-1 | CTX-M-14 | CTX-M-15 | CTX-M-27 | CMY-60 |
| A |  | 11 | 1 | 1 | 4 | 5 |  |
| B |  | 1 | 1 |  |  |  |  |
| C | C1-M27 | 23 |  |  |  | 23 |  |
|  | C1-nM27 | 5 |  | 3 |  | 2 |  |
|  | C2 | 54 |  |  | 54 |  |  |
| Unclassified |  | 7 |  |  | 4<sup>1</sup> | 3 | 1<sup>1</sup> |
| Total |  | 101 | 2 | 4 | 62<sup>1</sup> | 33 | 1<sup>1</sup> |

<sup>1</sup> One isolate belonging to an ST131 unclassified clade harboured both *bla*<sub>CTX-M-15</sub> and *bla*<sub>CMY-60</sub>.

##### Virotypes

ST131 has been reported in previous studies to be a highly virulent clone, exhibiting lethality in mouse sepsis models.^18, 31^ Virotypes of all 101 ST131 isolates were determined as previously described.^18, 32^ Virotype C was most common, represented by 38/101 (37.6%) of ST131 isolates and predominantly associated with *bla*_CTX-M-27_ (28/38 isolates) across clades A (n=3), C1-M27 (n=22), C1-nM27 (n=4), and one isolate belonged to an unclassified clade. All *bla*_CTX-M-27_-positive isolates belonged to virotype C, with the exception of one isolate for which a virotype could not be assigned. The association between virotype C and *bla*_CTX-M-27_ carriage is in agreement with a recent study conducted in France.^33^ Twenty-five isolates belonged to virotype A, all of which except one harboured *bla*_CTX-M-15_. The remaining 38 isolates belonged to either virotype B (n=1), D (n=1), G (n=8), or were unknown virotypes (n=28).

##### Genetic context of ESBLs/pAmpC genes

Despite the limitations of short read sequencing, by examining the contigs on which GOIs were located, we were able to determine the chromosomal or plasmid environments of ESBLs/pAmpC genes in 85/225 isolates. Forty CTX-M genes, of variants *bla*_CTX-M-14_ (n=3) and *bla*_CTX-M-15_ (n=37), were found to be located on the chromosome. These were found in 11 STs with STs 131 (n=11) and 73 (n=10) being the most represented. Whilst the CTX-M genetic environments were diverse in ST131, in ST73 9/10 isolates harboured the gene in the same genomic location, suggesting a high degree of clonality within this ST. The remaining GOIs were found to be located within a relatively diverse range of plasmids and across multiple STs. Interestingly all six DHA-1-harbouring isolates were found to harbour a similar IncI1 plasmid which was sequenced to closure during this study, pUB_DHA-1. Read mapping analyses showed that all six isolates exhibited 95-100% coverage and 98-100% identity against pUB_DHA-1.

##### Other important resistance genes found by WGS

Interestingly one ST69 CMY-2-producing isolate was also found to harbour *mcr-1*. Susceptibility testing, performed by broth microdilution, revealed that the MIC of colistin against this isolate is 8 mg/L and so it is colistin resistant. Attempts at transformation of *E. coli* DH5 alpha using plasmid DNA extracted from this isolate were successful indicating that *mcr-1* is plasmid encoded. Analysis of the genetic environment of *mcr-1* found that it is encoded on an IncI2 plasmid of approximately 62 kb and it lacks the upstream IS*Apl1* element that was described in the initial discovery of *mcr-1* in China.^34^ The plasmid itself does not encode any additional resistance genes and when subjected to NCBI BLAST analysis exhibited ∼96% similarity to *mcr-1* encoding plasmids found in both China (pHNGDF93; Genbank Accession Number MF978388) and Taiwan (p5CRE51-MCR-1; Genbank Accession Number CP021176) from animal and human origins, respectively. Since initial reports of its discovery in 2015,^34^ *mcr-1* has been reported worldwide in clinical *E. coli* isolates although remains relatively rare.

Another isolate, a *bla*_CTX-M-14_-positive ST38, also encoded the *bla*_OXA-48-like_ carbapenemase gene *bla*_OXA-244_.^35^ Transformation attempts using plasmid DNA from this isolate were unsuccessful and so it was concluded that *bla*_OXA-244_ is likely to be chromosomally encoded. Disc susceptibility testing showed that this isolate is resistant to ertapenem but susceptible to both imipenem and meropenem. The presence of the chromosomally-encoded OXA-48-like carbapenemase *bla*_OXA-244,_ confirms the observations of a previous study, where OXA-48-like genes were shown to have become embedded in the ST38 chromosome.^36^ ST38 is the most frequent ST associated with OXA-48-like enzymes in the UK.^36^

## Conclusions

Resistance to 3GCs and FQs in *E. coli* is of increasing concern due to the importance of these classes of drugs for the treatment of serious infections. As observed in this study and in line with previous reports globally,^2^ the dissemination of successful clones is a major cause of CTX-R in urinary isolates from primary care in South West England.

The correlation between CIP-R and CTX-R highlighted here can be largely attributed to the dominance of successful clones/clades, namely ST131 and ST1193; the majority of both harbour chromosomal FQ-R mutations. Through WGS of a subset of isolates we have shown that ST131 clade C2 is dominant and that the recently described ST131 subclade, C1-M27, is also prevalent despite not previously being described in the UK.

This study is the first analysis of CTX-R urinary *E. coli* derived from the community, and performed in a relatively localised area in South West England and can be useful for informing patient treatment as well as providing essential data for comparison purposes to other areas, both within and outwith the UK.

## Acknowledgements

Genome sequencing was provided by MicrobesNG (http://www.microbesng.uk), which is supported by the BBSRC (grant number BB/L024209/1). We are grateful to Aleksandra Pastuszek, Severn Infection Partnership, Southmead Hospital, for assistance in collecting the urinary *E. coli* isolates.

## Transparency declaration

None to declare.

## Supplementary data

Tables S1 and S2 are available as supplementary data.

## Supplementary Data

**Table S1.**
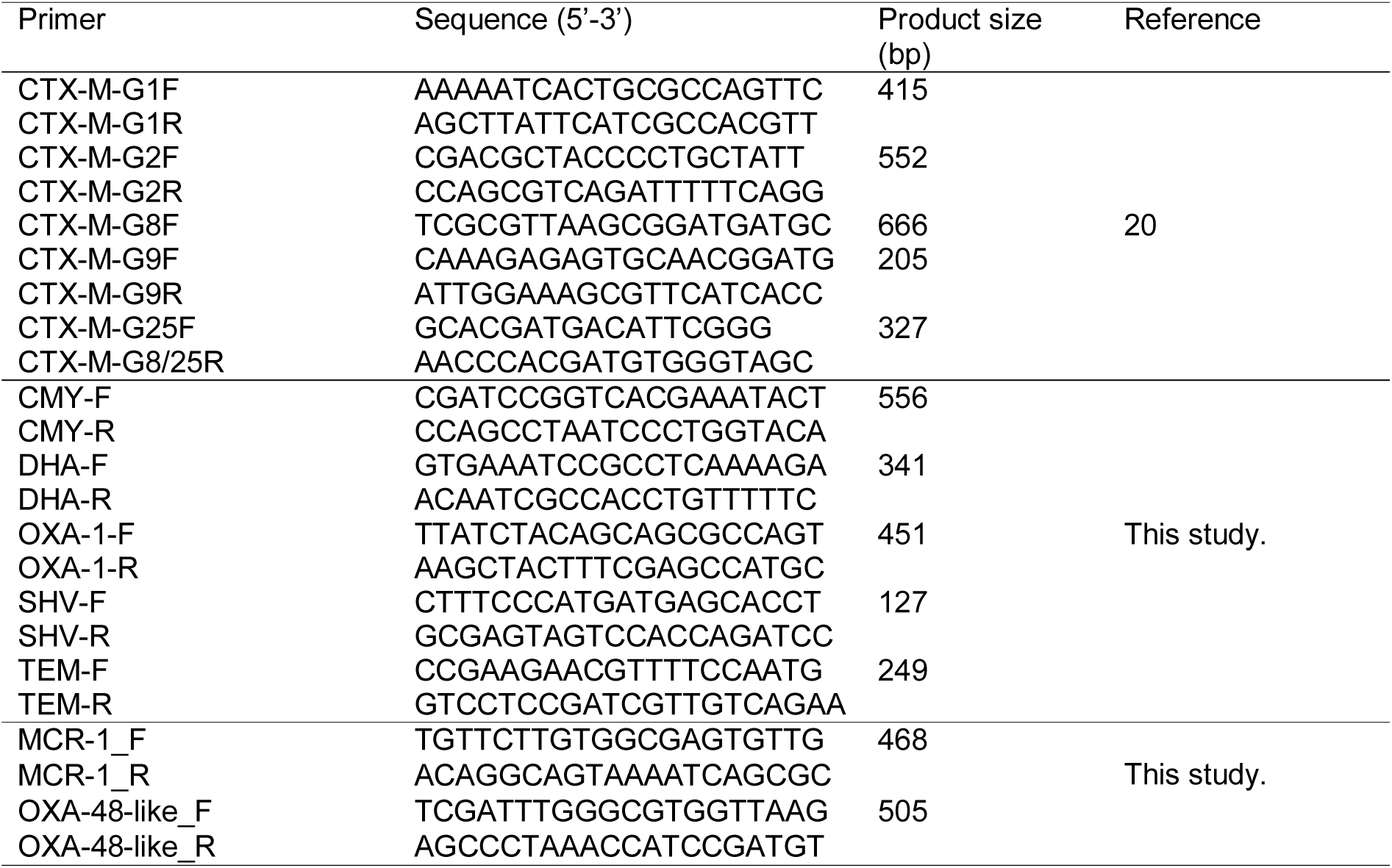
Primers used in this study.

**Table S2.** The STs and CTX-R mechanisms identified in 225 isolates subjected to WGS.

| STs<br>(no. if >1) | Beta-lactamase GOIs/mechanism identified (no.) |
| --- | --- |
| 131 (101) | CTX-M-15 (62), CTX-M-27 (33), CTX-M-14 (4), CTX-M-1 (2), CMY-60<sup>1</sup> |
| 69 (15) | CTX-M-15 (9), CTX-M-1, CTX-M-27, CTX-M-55, DHA-1<sup>2</sup>, CMY-2 (2), CMY-42, CMY-60<sup>3</sup> |
| 73 (15) | CTX-M-15 (12), AmpC-hyp (3) |
| 38 (13) | CTX-M-15 (5), CTX-M-27 (4), CTX-M-14 (3), DHA-1, OXA-244<sup>4</sup> |
| 1193 (11) | CTX-M-27 (5), CTX-M-15 (3), CMY-2 (2), CTX-M-55 |
| 10 (8) | CTX-M-15 (6), CTX-M-55, CMY-60<sup>5</sup>, CTX-M-14 |
| 359 (4) | SHV-12 (4) |
| 141 (3) | CTX-M-1 (2), AmpC-hyp |
| 200 (3) | AmpC-hyp (3) |
| 209 (3) | CTX-M-3 (3) |
| 349 (3) | DHA-1 (2), CTX-M-15 |
| 394 (3) | CTX-M-15 (3) |
| 636 (3) | CTX-M-15 (3) |
| 12 (2) | CTX-M-14, SHV-12 |
| 58 (2) | CTX-M-14, CTX-M-55 |
| 80 (2) | CTX-M-15, CMY-2 |
| 127 (2) | CTX-M-1, DHA-1 |
| 405 (2) | CTX-M-15 (2) |
| 648 (2) | CTX-M-15, CMY-42 |
| 963 (2) | CMY-2 (2) |
| 1722 (2) | CTX-M-1, CTX-M-27 |
| 8313 (2) | CTX-M-15 (2) |
| 23 | CTX-M-1 |
| 54 | AmpC-hyp |
| 75 | AmpC-hyp |
| 88 | AmpC-hyp |
| 95 | CTX-M-15 |
| 155 | AmpC-hyp |
| 162 | SHV-12 |
| 224 | CTX-M-15 |
| 428 | AmpC-hyp |
| 443 | CTX-M-15 |
| 448 | AmpC-hyp |
| 457 | CMY-4 |
| 491 | DHA-1 |
| 540 | CTX-M-8 |
| 1431 | CTX-M-15 |
| 2015 | CTX-M-1 |
| 2914 | CTX-M-15 |
| 3036 | DHA-1 |
| 4981 | CTX-M-15 |
| 7401 | DHA-1 |
| 8312 | CTX-M-15 |
| 8467 | CTX-M-15 |

<sup>1</sup> – CMY-60 harboured alongside CTX-M-15.
<sup>2</sup> – DHA-1 harboured alongside CTX-M-15.
<sup>3</sup> – CMY-60 harboured alongside CTX-M-15.
<sup>4</sup> – OXA-244 harboured alongside CTX-M-14.
<sup>5</sup> – CMY-60 harboured alongside CTX-M-15.

